# Chloroplast genome engineering of potato enables diterpene production without agronomic penalty

**DOI:** 10.64898/2026.05.15.725540

**Authors:** Alessandro Occhialini, Xinlu Chen, Samantha A. Miller, Mohammad Majdi, Ivette A. Fuentes Quispe, Gabriella King, Feng Chen

## Abstract

Terpenes constitute the largest and most structurally diverse class of plant secondary metabolites, with critical roles in plant-environment interactions and broad industrial applications. Although nuclear genome engineering of terpene pathways has been extensively explored, chloroplast genome engineering remains largely undeveloped, with all reported studies restricted to the model plant *Nicotiana*. Here we report successful chloroplast genome engineering for diterpene production in the crop plant potato (*Solanum tuberosum*). First we identified the *trnT/trnL* plastomic locus as optimal for minimizing integration-associated growth penalties. Insertion of a bifunctional diterpene synthase gene into this plastomic site yielded transplastomic plants with successful diterpene production, but with reduced growth. The co-expression of a geranylgeranyl diphosphate synthase gene to enhance precursor supply restored normal growth while elevating diterpene accumulation. Transplastomic plants were otherwise agronomically comparable to wild-type. This work expands chloroplast engineering as a viable strategy for terpene pathway engineering in crop improvement and high-value terpene production.

## INTRODUCTION

Terpenes represent the largest and most structurally diverse class of plant secondary metabolites ^1^. They serve critical biological roles in plants, including defense against herbivores and pathogens, and attraction of pollinators ^1,2^. Owing to their broad biological activities, numerous terpenes have been developed as pharmaceuticals, including the antimalarial artemisinin and the anticancer agent Taxol ^3–5^. Their high energy density has also made terpenes attractive candidates as next-generation biofuels ^6,7^. The combination of ecological importance and wide-ranging applications has driven substantial interest in enhancing terpene biosynthesis and developing scalable production platforms. Two major production platforms have been explored. The first is microbial-based systems, including bacteria and yeast, which offer well-established genetic tools and rapid design-build-test cycles ^8–10^. Significant progress has been made in engineering microbial hosts for terpene production, and some microbial-derived terpenes have reached commercial scale ^11,12^. The second platform is plant-based production, which offers distinct advantages in terms of sustainability, use of solar energy, and access to native biosynthetic machinery and precursor pools ^13–15^. However, despite significant advances, progress in plant-based terpene engineering has lagged behind microbial systems, largely due to the complexity of plant genomes, limited transformation efficiencies in many crop species, and the challenge of achieving high product yields without compromising plant growth and agronomic performance.

Unlike microbial systems, terpene biosynthesis in plants is compartmentalized between the cytosol, where the mevalonate (MVA) pathway generates sesquiterpene and triterpene precursors, and the plastid, where the methylerythritol phosphate (MEP) pathway supplies precursors for monoterpenes, diterpenes, and tetraterpenes ^16–18^. Much work has been done with nuclear genome engineering of terpene pathways across diverse plant species, achieving significant gains in terpene accumulation ^19–21^. However, an alternative and potentially more powerful strategy is to engineer the chloroplast genome directly. Chloroplast genome engineering offers several compelling advantages, including high transgene copy number due to organelle polyploidy, the capacity to express multiple genes in operons facilitating metabolic engineering, absence of epigenetic gene silencing, and transgene biocontainment through maternal inheritance ^22–26^. Crucially, the plastid stroma harbors the native MEP pathway ^27,28^, providing a metabolically primed environment rich in terpene precursors into which heterologous terpene biosynthesis genes can be inserted. Despite these advantages, reports of chloroplast genome engineering for the terpene pathway remain remarkably less explored. Studies to date have included enhancement of the MEP pathway via the overexpression of 1-deoxy- D-xylulose 5-phosphate reductoisomerase gene ^29^, introduction of a non-native carotenoid branch pathway ^30^, insertion of the complete yeast MVA pathway to boost precursor supply ^31^, engineering of the artemisinin pathway using multi-gene stacking and hybrid compartmentalization strategies ^32^, squalene biosynthesis via farnesyl diphosphate synthase (FPS) and a squalene synthase (SQS) insertion ^33^, and most recently, introduction of taxadiene synthase for Taxol precursor production ^34^. Critically, all of these studies were conducted exclusively in the model plant *Nicotiana tabacum*, leaving it entirely unknown whether chloroplast genome engineering represents a viable strategy for terpene production in other plant species, particularly crop plants of agricultural importance.

In this study, our goal is to expand diterpene pathway engineering via chloroplast genome modification in a food crop plant. We selected potato (*Solanum tuberosum*) as our target species, given its global agricultural importance as one of the world’s most widely grown food crops ^35^ and its well-established status as a plant system for chloroplast genome engineering ^36,37^. This study pursues two key novelties. First, we applied an evolution-guided design strategy for terpene pathway construction. Recent phylogenomic studies on the evolution of isoprenoid diphosphate synthase (*IDS*) genes have demonstrated that geranylgeranyl diphosphate (GGPP) synthase (*GGPPS*) genes in glaucophytes, the basal lineage of the plant kingdom, remain encoded by the plastome ^38^, providing an evolutionary rationale for introducing such *GGPPS* gene directly into the chloroplast genome of potato. Furthermore, comprehensive evolutionary analyses of terpene synthases (TPS), which are pivotal for terpene biosynthesis ^39–41^, have shown that class I&II bifunctional diterpene synthases (diTPS) are absent in flowering plants but present across all other land plant lineages ^18,42,43^, suggesting secondary gene loss in angiosperms. Accordingly, we selected a bifunctional diTPS from a non-flowering plant lineage to mimic the evolutionary restoration of this biosynthetic capacity in potato. Second, to avoid the potential growth penalties associated with transgene integration, we identified and characterized a new integration site within the potato plastome as optimal for minimizing integration-associated growth penalties. This new integration site was used for the diterpene pathway engineering producing transplastomic potato plants without any phenotypic penalties compared to wild-type control plants. Collectively, this study demonstrates that chloroplast genome engineering can serve as a powerful platform for evolution-guided terpene pathway engineering in crop plants.

## RESULTS

### Development of an optimal plastomic integration site for transgene integration

One of the major limitations that has prevented the use of transplastomic plants in agriculture is the frequent occurrence of phenotypic penalties caused by several intrinsic factors, including excessive transgene expression, overloading of endogenous metabolic pathways, and negative effects of the transgene cassette on endogenous genetic elements at the integration site ^44–46^. In this study, to develop a transgene integration site with reduced or no impact on plant phenotype due to potential perturbation of endogenous genetic elements, a chloroplast transformation vector for transgene integration into the *trnT*/*trnL* site of the large single copy (LSC) region of potato plastome was tested. In fact, this transformation vector induces transgene integration in an intergenic region of ∼0.7 kb in between upstream (*rps4* and *trnT*) and downstream (*trnL* and *trnF*) genes with transcription orientation at opposite direction of the integration site (Figure 1A) ^47^, representing a potential optimal integration site for transgene integration with a minimum perturbation of endogenous genetic elements.

**Figure 1:**
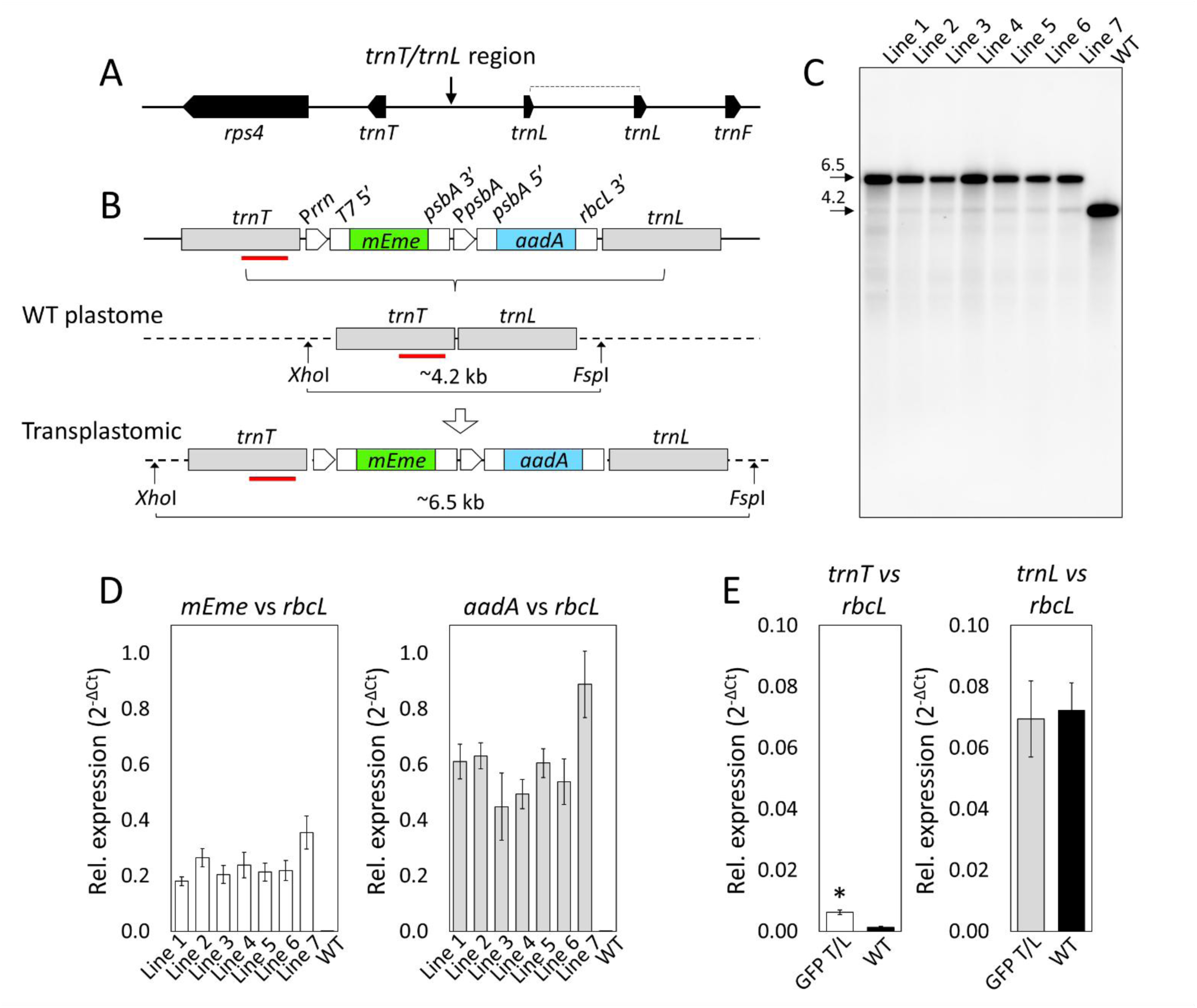
Genotyping of GFP T/L transplastomic lines. **(A)** Schematic representation of the 3 kb plastomic region surrounding the *trnT*/*trnL* transgene integration site (arrow). The *rps4*, *trnT*, *trnL*, *trnF* genes are indicated. **(B)** Schematic representation of the GFP T/L vector rearrangement into the *trnT/trnL* integration site of potato plastome. The GFP T/L vector contains an *mEmerald* (green fluorescent reporter gene) and an *aadA* (spectinomycin resistance gene) expression cassettes in between ∼1.5 kb left and right arms (gray) homologous to the *trnT/trnL* site. The location of a ∼0.5 kb-long probe specific to the left arm is indicated. The molecular weights of DNA fragments obtained using the restriction enzyme combination *Xho*I/*Fsp*I is also indicated. Regulatory elements for expression of *mEmerald* (*mEme*) and *aadA* (spectinomycin resistance gene) are indicated: *N. tabacum* P*rrn-T7G10* promoter-5’UTR fusion (P*rrn* and T75’); *N. tabacum psbA* 3’UTR (*psbA3’*); *N. tabacum* promoter-5’UTR of *psbA* (P*psbA* and *psbA5’*); and *N. tabacum rbcL* 3’UTR (*rbcL3’*). **(C)** Southern blot performed using the ∼0.5 kb probe and genomic DNA preparations from leaves of GFP T/L lines 1-7. A DNA sample from wild-type potato plant is used as negative control. Molecular weights of DNA bands (kb) are indicated in the blots. **(D)** qRT-PCRs were performed using cDNA samples from GFP T/L plants along with wild-type controls. Primers for detection of *mEmerald* and *aadA* transgenes have been used. Graphs represent the relative expression (2^-ΔCt^) of the indicated gene versus the plastome reference gene *rbcL*. **(E)** qRT-PCRs were performed using cDNA samples from GFP T/L and wild-type controls, and primers for amplification of *trnT* and *trnL* located in proximity of the integration site. Graphs represent the relative expression (2^-ΔCt^) of the indicated gene versus the *rbcL* gene expressed as the average of all GFP T/L lines (1-7). For D and E, data represent mean ± SE (standard error) of 3 biological replicates per each line and 3 technical replicates per each biological. Mean separation was evaluated using ANOVA Post-Hoc Dunnett’s T3 (p<0.05) (D) and t test (p<0.05) (E).

The T/L transformation vector was designed with a transgene cassette for expression of the *mEmerald* reporter gene (green-fluorescent reporter) and the selectable marker gene *aadA* (spectinomycin resistance) cloned in between 1.5-kb-long homologous arms (left and right) designed on the plastomic *trnT*/*trnL* region (Figure 1A). In this GFP T/L transformation vector, the *mEmerald* expression was under the control of the *N. tabacum rrn S16* gene promoter fused to the *T7G10* leader and the 3’UTR from *N. tabacum psbA* gene, while the *aadA* expression was under the control of the promoter-5’UTR from the *N. tabacum psbA* gene and the 3’UTR from the *N. tabacum rbcL* gene (Figure 1B). The GFP T/L vector was delivered into potato leaf disks by biolistic and seven independent transplastomic plants were produced in tissue culture. Southern blots performed using a ∼0.5 kb probe designed against the *rps4* gene located in the left arm confirmed correct transgene cassette integration into the *trnT*/*trnL* site of the plastome. The presence of a major DNA band at ∼6.5 kb indicates *mEmerald*-*aadA* cassette integration in prevalence at homoplasmy into the *trnT*/*trnL* site of all GFP T/L lines (Figure 1C). In order to support correct expression of transgenes in transplastomic lines, qRT-PCRs were performed using cDNA samples obtained from leaf tissue and oligonucleotides specific to the two transgenes (*mEmerald* and *aadA*). While all lines were statistical identical for expression of both transgenes, the expression levels of *mEmerald* and *aadA* were different, corresponding to the ∼24% and ∼60% of the expression level of the endogenous *rbcL* reference gene, respectively (Figure 1D). Moreover, integration of the *mEmerald*-*aadA* cassette into the *trnT*/*trnL* site of plastome had minimal impact on the expression of genes at this integration site. In fact, qRT-PCR analysis indicates that the downstream *trnL* gene expression is unaffected, while the upstream *trnL* gene is only minimally affected compared to the wild-type unmodified locus (Figure 1E).

### Transgene integration into the *trnT*/*trnL* site of the plastome had no negative impact on plant growth and phenotype

To verify that transgene cassette integration into the *trnT*/*trnL* site of plastome was not associated with any phenotypic alterations, a growth experiment was performed in a greenhouse space. For this purpose, all GFP T/L lines along with wild-type control plants were grown until the appearance of the first floral bolt (anthesis), and then, several phenotypic paraments were evaluated (Figure 2). In this study, lines 2, 3 and 5 showed the same plant height compared to wild-type plants, while lines 1, 4, 6 and 7 appeared to have an overage of ∼27% reduction in plant height (Figure 2B). On the contrary, all GFP T/L lines showed the same or higher ability to accumulate total biomass (fresh and dry) per unit of stem height, foliar biomass (fresh and dry) per unit of leaf area, and leaf chlorophyll content compared to wild-type control plants (Figure 2C-G). While phenotypic variation was observed for transplastomic lines obtained with the same construct, these results indicate that selected GFP T/L lines 2, 3 and 5 can reach the same phenotypic parameters compared to wild-type control plants, demonstrating that transgene integration into *trnT*/*trnL* site of plastome is not associated with any phenotypic alterations.

**Figure 2:**
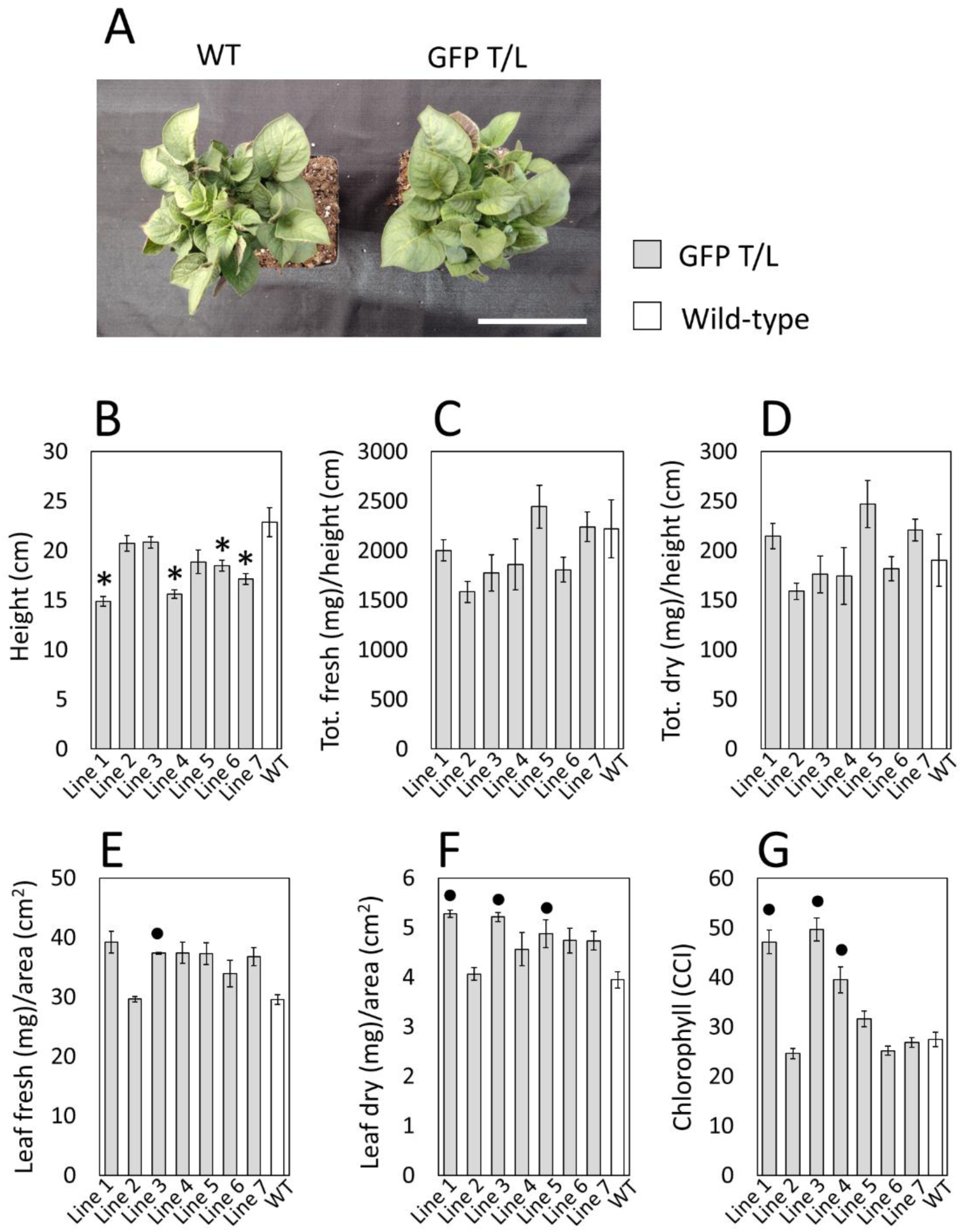
Growth characteristics of GFP T/L transplastomic lines. **(A)** Images showing ∼5-6-week-old GFP T/L lines (1-7) along with wild-type (WT) control plants grown on pot in a greenhouse space. Scale bar: 10 cm. Various phenotypic plant characteristics collected at anthesis ∼7-8-week-old are shown: **(B)** Plant height (cm); **(C)** ratio of total fresh weight (mg) to plant height (cm); **(D)** ratio of total dry weight (mg) to plant height (cm); **(E)** ratio of leaf fresh weight (mg) to foliar area (cm^2^); **(F)** ratio of leaf dry weight (mg) to foliar area (cm^2^); **(G)** chlorophyll content index (CCI). Data are expressed as mean ± standard error (SE) of 4 plants per each transplastomic line and wild-type plants. Means were compared using ANOVA Post-Hoc Dunnett’s t test (p<0.05, B-D and F) or Post-Hoc Dunnett’s T3 (p<0.05, E and G) and statistical significance lower (*) or higher (•) compared to the wild-type reference group are indicated.

As we described in previous investigations, phenotypic variation can also be observed for transplastomic potato plants produced with the same site-specific transgene integration vector ^48,49^, justifying the requirement of characterizing different independent lines for a comprehensive phenotypic analysis. While all transplastomic lines produced with the same vectors are genetically identical for the location of transgene integration, it is likely that phenotypic variation is due to unpredicted genetic and epigenetic alterations, known as somaclonal variation, frequently associated with procedures of plant regeneration and maintenance in tissue culture ^50–52^.

### Transplastomic potato plants with a bifunctional diterpene synthase produced diterpenes but showed severe phenotypic penalties

Plants employ the cytosolic MVA pathway to produce farnesyl diphosphate for sesquiterpene biosynthesis and the plastidic MEP pathway to produce GGPP and geranyl diphosphate (GPP) for the biosynthesis of diterpenes and monoterpenes, respectively ^16–18^. Whereas the production of GPP has evolved multiple times during land plant evolution ^53^, the production of GGPP and diterpenes in plastids using nuclear genes is conserved across land plants ^16,18,54^. As such, diterpene synthase enzymes encoded by the plastome could naturally access the GGPP pool within plastids. Therefore, we aimed in this study to engineer the diterpene biosynthetic pathway in potato through chloroplast genome engineering.

Diterpene synthases belong to the TPS family, which is a mid-sized family among land plants. TPSs contain seven subfamilies: TPS-a, -b, -c, -d, -e/f, -g, and -h. Diterpene synthases generally belong to the TPS-c and TPS-e/f subfamilies. Our analysis of the sequenced potato genome showed that potato contains 33 *TPS* genes encoding proteins longer than 400 amino acids with 3 members in the TPS-c subfamily and 3 members in the TPS-e/f subfamily. All members of the TPS-c subfamily contain only the DxDD motif, signature of class I diterpene synthase, while all members of the TPS-e/f subfamily of potato contain the DDxxD motif, signature of class II diterpene synthase (Figure S1). It is known that class I&II bifunctional diterpene synthases occur in non-flowering plants ^18^. Class I&II bifunctional enzymes catalyze sequential reactions and therefore can control product specificity and may have less impact on natural biosynthetic pathways. Therefore, we chose PdTPS, a class I&II diterpene synthase from the lycophyte *Phylloglossum drummondii* ^18^ as our target transgene. PdTPS had been previously demonstrated to catalyze the formation of levopimaradiene from GGPP using an engineered bacterial system ^18^. To fully characterize this enzyme, recombinant PdTPS produced in *E. coli* was analyzed in *in vitro* enzyme assays using GGPP as substrate. In addition to levopimaradiene, a second diterpene, dehydroabietane, was produced (Figure 3A and S2).

**Figure 3.**
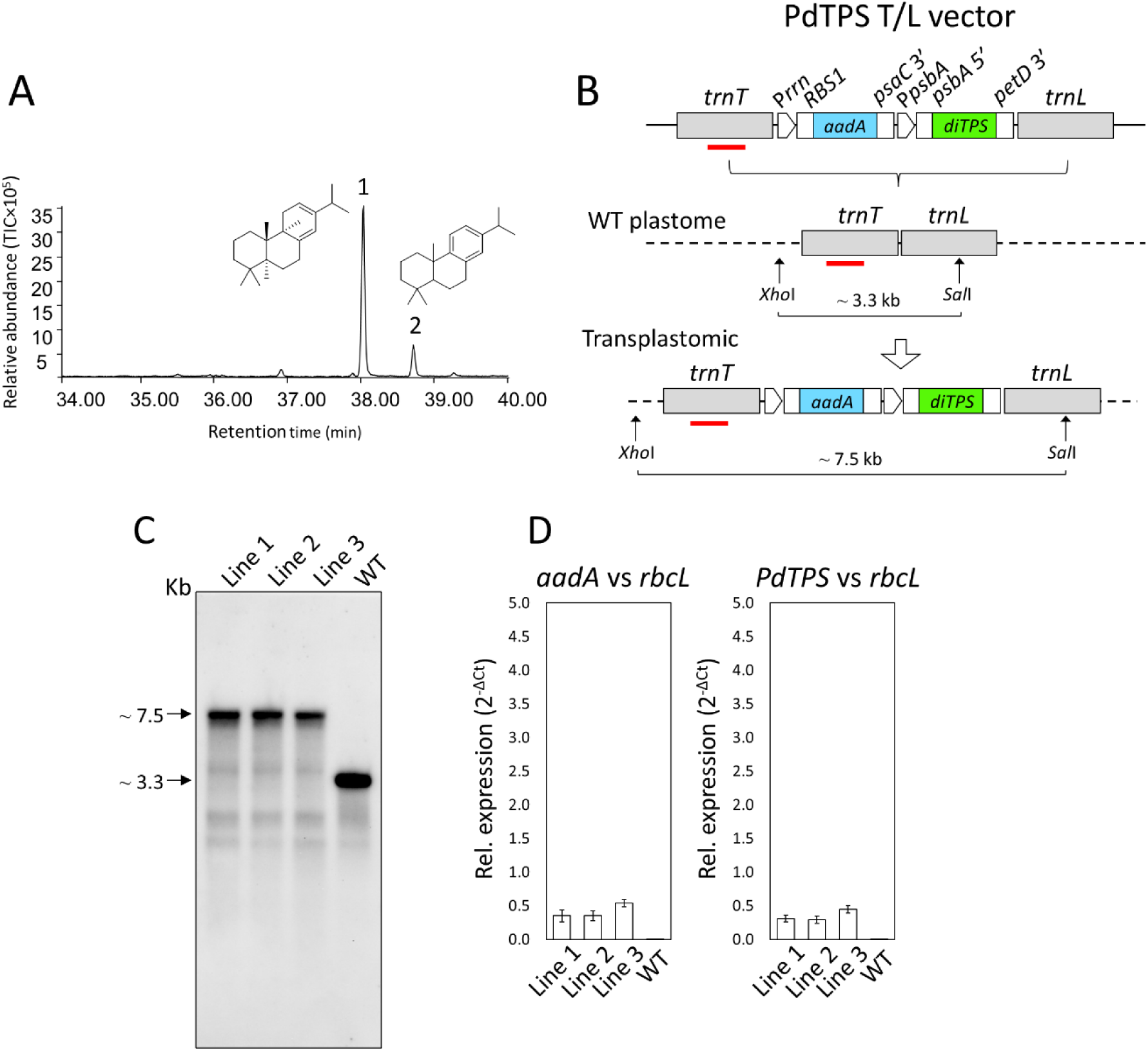
Genotyping and molecular characterization of PdTPS T/L transplastomic lines. **(A)** GC chromograms of *in vitro* enzymatic assay using PdTPS. Graphs of the relative abundance of terpenoids (TICx10^5^) versus the retention time (minutes) are indicated. The two peaks correspond to levopimaradiene (1) and phenanthrene (2), respectively. The molecular structures of the two terpenes are also indicated. **(B)** Schematic representation of PdTPS T/L vector rearrangement into the *trnT/trnL* integration site of potato plastome. The PdTPS T/L vector contains the *aadA* (spectinomycin resistance gene) and the *PdTPS* (diterpene synthase) expression cassettes cloned between ∼1.5 kb left and right arms (gray) homologous to the *trnT/trnL* integration site. Regulatory elements for expression of *aadA* (spectinomycin resistance) and *PdTPS* (diterpene synthase) are indicated: *N. tabacum rrn16* promoter (P*rrn*) fused to the ribosome binding site (*RBS1*) of the *rbcL* 5’UTR; *N. tabacum psaC* gene 3’UTR (*psaC* 3’); *N. tabacum* promoter-5’UTR of *psbA* (P*psbA* and *psbA* 5’); *N. tabacum petD* gene 3’UTR (*petD* 3’). The location of 0.5 kb-long probe is indicated with a red bar. The molecular weights of DNA fragments obtained using the restriction enzyme combination *Xho*I/*Sal*I are indicated. **(C)** Southern blot performed using a 0.5 kb probe complementary to the *trnT* arm (*rps4* gene) and genomic DNA preparations from leaves of PdTPS T/L (1-3) lines along with wild-type control plants. Molecular weights of DNA bands (kb) are indicated in the blots. **(D)** qRT-PCRs performed using cDNA samples obtained from PdTPS T/L lines (1-3) and wild-type control plants, and specific primers for detection of *aadA* and *PdTPS* transgenes. Graphs represent the relative expression (2^-ΔCt^) of the indicated transgene vs the plastome reference gene *rbcL*. Data represent mean ± SE (standard error) of 3 biological replicates per each line and 3 technical replicates per each biological. Mean separation was evaluated using ANOVA Post-Hoc Dunnett’s T3 (p<0.05).

To proceed with diterpene pathway engineering in potato, a PdTPS T/L transformation vector, for transgene integration into the *trnT*/*trnL* site of plastome was produced (Figure 3B). As shown in Figure 2, the expression of dicistronic expression cassette into the *trnT*/*trnL* site was not associated with plant phenotypic penalties, representing an optimal choice for producing transplastomic plants to use in agriculture for improving traits. To provide appropriate transgene expression in plant tissues, strong endogenous regulatory elements were used. In this vector the expression of the *aadA* marker-gene was under the control of the *N. tabacum rrn S16* promoter fused to the ribosome binding site (RBS) of the *rbcL* gene at 5’UTR and the *N. tabacum psaC* gene terminator at the 3’UTR, while the *PdTPS* expression was under the control of the *N. tabacum* promoter-5’UTR of the *psbA* gene and the 3’UTR of the *N. tabacum petD* gene (*petD* 3’) (Figure 3B). The PdTPS T/L vector was delivered into potato leaf disks and a total of three independent transplastomic lines (PdTPS lines 1-3) were produced in tissue culture. Southern blot analysis performed using DNA genomic preparations from leaf tissue and a ∼0.5 kb probe complementary to the *trnT* left arm (*rps4* gene) confirmed correct transgene integration at homoplasmy in all transplastomic lines (Figure 3C). In fact, correct PdTPS T/L vector integration was confirmed by the presence of ∼7.5 kb bands at the predicted molecular weight, while the absence of a ∼3.3 kb band corresponding to the unmodified wild-type *trnT*/*trnL* integration site confirmed transgene cassette integration at homoplasmy (Figure 3C).

In order to support correct expression of transgenes in PdTPS T/L lines, qRT-PCRs were performed using cDNA samples obtained from leaf tissue and oligonucleotides specific to *aadA* and *PdTPS*. These results confirmed correct expression of all transgenes and that the expression of individual transgenes was not statistically different in transplastomic lines produced with the same construct (Figure 3D). A minimum difference of expression levels of *aadA* and *PdTPS* was observed, corresponding to the ∼40% and ∼35% of the expression level of the endogenous *rbcL* gene. Transgene cassette integration into *trnT*/*trnL* plastomic site had also no impact on the expression of *trnT* and *trnL* genes compared to the wild-type control (Figure S3A and B), supporting previous qRT-PCR results obtained in GFP T/L lines (Figure 1E). Next, leaves from 4-week-old PdTPS-transplastomic potato plants in tissue culture were subjected to metabolic profiling using gas chromatography-mass spectrometry (GC-MS). While no diterpenes were detected from wild-type potato plants at the same developmental stage, two diterpenes, levopimaradiene and dehydroabietane, were detected from transplastomic plants (Figure 4A and B). However, in tissue culture, PdTPS T/L lines showed serious phenotypic penalties including small plant size, bushy phenotype, and reduced rooting (Figure S4).

**Figure 4.**
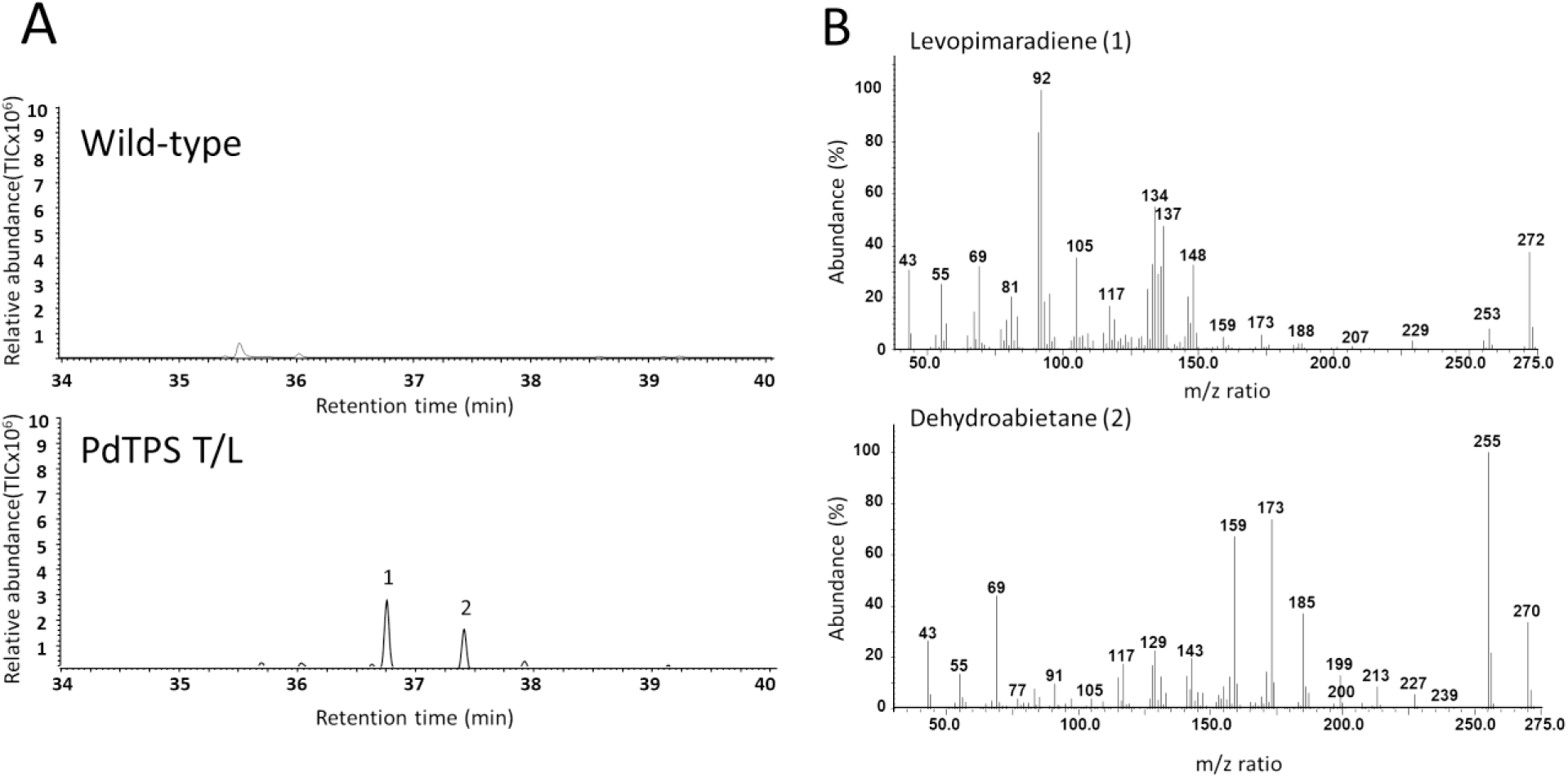
GC/MS analysis of terpene production in PdTPS T/L transplastomic lines. **(A)** GC chromograms of metabolic analysis of PdTPS T/L transplastomic lines and wild type control plants. Graphs of the relative abundance of terpenes (TICx10^5^) versus the retention time (minutes) are indicated. The two peaks correspond to levopimaradiene (1) and dehydroabietane (2), respectively. **(B)** Mass spectrums of levopimaradiene (1) and dehydroabietane (2) from PdTPS T/L transplastomic lines. The relative abundance (%) and the m/z ratio are indicated in the y and x axis, respectively.

### The expression of a geranylgeranyl diphosphate synthase along with the bifunctional diterpene synthase induces a higher accumulation of diterpenes and restores a normal plant phenotype

GGPP, the substrate for diterpene biosynthesis, also serves as a precursor for multiple essential plant metabolites, including chlorophyll, carotenoids, and the phytohormone gibberellins ^55–57^. In engineered potato plants, diversion of GGPP toward the production of levopimaradiene and dehydroabietane is expected to reduce metabolic flux into these native pathways. Consequently, this reallocation may negatively impact plant physiology, growth, and development, as we observed (Figure S4). To mitigate these negative effects, it is important to increase the GGPP pool within plastids. As mentioned earlier, GGPP is synthesized by GGPPS using universal isoprenoid precursors derived from the MEP pathway ^16–18^. Given the central role of GGPPS in GGPP biosynthesis, a logical strategy to enhance GGPP availability is to increase GGPPS expression levels. GGPPS is a highly conserved enzyme across the plant kingdom. A recent study suggests that plant GGPPS originated from cyanobacteria through endosymbiosis ^38^. Notably, in glaucophytes, an early-diverging lineage of the plant kingdom, the GGPPS gene remains encoded in the plastome ^38^. We reason that such plastid-encoded enzymes may be particularly suitable for chloroplast genome engineering. Therefore, we selected the *GGPPS* gene from the glaucophyte *Cyanophora biloba*, designated as *CbGGPPS*, as our target transgene.

For production of transplastomic plants, a chloroplast transformation vector (PdTPS-CbGGPPS T/L vector), for *aadA-PdTPS-CbGGPPS* cassette integration into the *trnT*/*trnL* site of plastome was designed (Figure 5A). In this vector the expression of the *aadA* selectable marker and the *PdTPS* transgene was under the control of the same regulatory elements (promoters and UTRs) described for the PdTPS T/L vector, while the downstream *CbGGPPS* transgene was under the control of the *N. tabacum rbcL* promoter fused to a synthetic ribosome binding site and the 3’UTR of the *Lactuca sativa psbA* gene (Figure 5A). A total of six independent transplastomic lines with normal phenotype were produced in tissue culture and genotyped by Southern blots using a ∼0.5 kb probe complementary to *trnT* left arm. The presence of only one band at the predicted molecular weight for *aadA-PdTPS-CbGGPPS* cassette integration (∼8.9 kb) confirmed homoplasmy of all transplastomic lines (Figure 5B).

**Figure 5.**
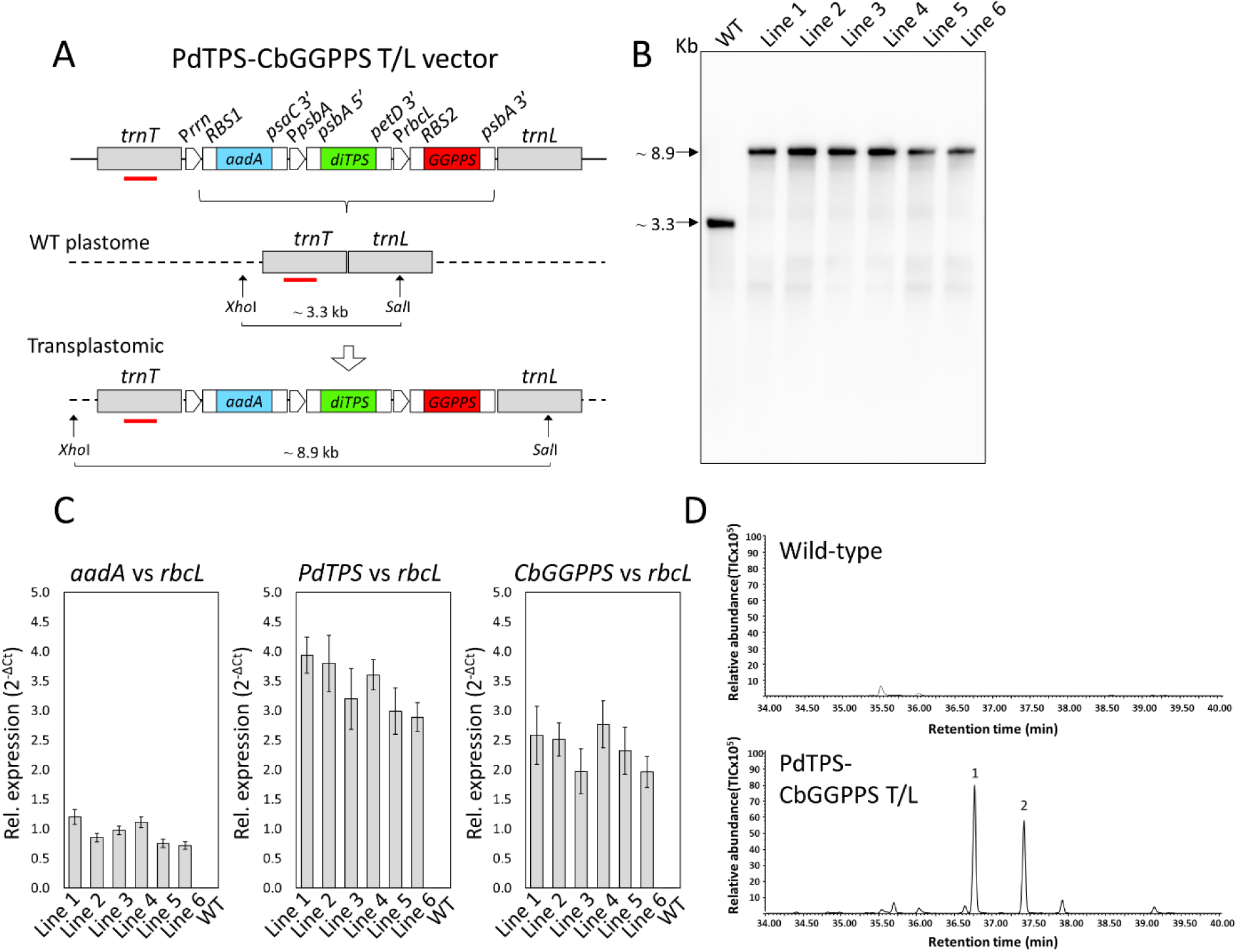
Genotyping and molecular characterization of PdTPS-CbGGPPS T/L transplastomic lines. **(A)** Schematic representation of PdTPS-CbGGPPS T/L vector rearrangement into the *trnT/trnL* integration site of potato plastome. The PdTPS-CbGGPPS T/L vector contains the *aadA* selectable marker (spectinomycin resistance gene) along with the *PdTPS* (diterpene synthase) and the *CbGGPPS* (geranylgeranyl pyrophosphate synthase) gene expression cassettes inserted between ∼1.5 kb left and right arms (gray) homologous to the *trnT/trnL* integration site. Regulatory elements for transgene expression are indicated: *N. tabacum rrn16* promoter (P*rrn*) fused to the ribosome binding site (*RBS1*) of the *rbcL* 5’UTR; *N. tabacum psaC* gene 3’UTR (*psaC* 3’); *N. tabacum* promoter-5’UTR of *psbA* (P*psbA* and *psbA* 5’); *N. tabacum petD* gene 3’UTR (*petD* 3’); *N. tabacum rbcL* promoter fused to a synthetic ribosome binding site (*RBS2*); and the *Lactuca sativa psbA* gene 3’UTR (*psbA* 3’). The location of a 0.5 kb probe is indicated with a red bar. The molecular weights of DNA fragments obtained using the restriction enzyme combination *Xho*I/*Sal*I are indicated. **(B)** Southern blot performed using a 0.5 kb probe complementary to the *trnT* arm (*rps4* gene) and genomic DNA preparations obtained from PdTPS-CbGGPPS T/L (1-6) lines, and wild-type controls. Molecular weights of DNA bands (kb) are indicated in the blots. **(C)** qRT-PCRs performed using cDNA samples obtained from PdTPS-CbGGPPS T/L lines (1-6) and wild-type control plants, and specific primers for detection of *aadA*, *PdTPS*, and *CbGGPPS* transgenes. Graphs represent the relative expression (2^-ΔCt^) of the indicated transgene vs the plastome reference gene *rbcL*. Data represent mean ± SE (standard error) of 3 biological replicates per each line and 3 technical replicates per each biological. Mean separation was evaluated using ANOVA Post-Hoc Dunnett’s T3 (p<0.05). **(D)** GC chromograms of metabolic analysis of PdTPS-CbGGPPS T/L transplastomic lines and wild type control plants. Graphs of the relative abundance of terpenes (TICx10^5^) versus the retention time (minutes) are indicated. The two peaks correspond to levopimaradiene (1) and dehydroabietane (2), respectively. The graph of wild-type control is the same as Figure 4A, since the experiments were performed at the same time.

In the next step, correct expression of the three transgenes (*aadA*, *PdTPS* and *CbGGPPS*) was confirmed by qRT-PCR. Also in this case, transgene expression levels were not statistically different in transplastomic lines produced with the same construct (Figure 5C). On the contrary, different transgene expression levels were observed comparing this *PdTPS-CbGGPPS* genotype with previously described lines expressing only *PdTPS*. As indicated before, while in PdTPS T/L lines the expression of *aadA* and *PdTPS* was the ∼40% and ∼35% of the expression level of the endogenous *rbcL* gene (Figure 3D), in PdTPS-CbGGPPS T/L lines the *aadA* expression was nearly the same as *rbcL* whereas the expression of *PdTPS* and *CbGGPPS* was ∼2.4 and ∼1.4-fold higher than *rbcL*, respectively (Figure 5C). These differences in expression levels could be due to a different construct design (two versus three transgene cassettes) and the presence of particularly strong regulatory elements in the third *CbGGPPS* cassette, such as the lettuce *psbA* 3’UTR ^58,59^, that could improve stability of the polycistronic transcript transcribed from the upstream P*rrn* promoter. Despite these higher *aadA*, *PdTPS* and *CbGGPPS* gene expressions, also in PdTPS-CbGGPPS T/L plants, transgene cassette integration had no impact on the expression levels of *trnT* and *trnL* genes of the plastomic site (Figure S5A and B). In accordance with the higher transgene expression and with the presence of an extra GGPP pool of precursors, PdTPS-CbGGPPS T/L lines produced ∼2.5-fold the amount of diterpenes compared to PdTPS T/L plants based on peak area comparison (Figure 5D).

### Transplastomic potato plants producing diterpenes showed normal growth and yield

While the genetic engineering toolbox for chloroplasts holds great potential for improving crop traits, its application in agriculture remains limited. Although chloroplasts have been successfully used as bioreactors to produce valuable molecules, transplastomic approaches have found limited application to enhance crop performance, mainly due to the frequent phenotypic penalties associated with chloroplast genome modification in transplastomic plants. Therefore, in the following fundamental step of this study, to verify that the heterologous production of diterpenes was not associated with any phenotypic penalties, PdTPS-CbGGPPS T/L lines and wild-type control plants were analyzed in a growth experiment.

For this purpose, transplastomic plants and wild-type controls were grown until the appearance of the first floral bolt (anthesis), and various phenotypic parameters were evaluated (Figure 6). At this developmental stage, the plant height of all PdTPS-CbGGPPS T/L lines was not statistically different compared to wild-type plants (Figure 6B). Among these transplastomic plants, lines 4-6 were also not statistically different compared to wild-type plants for their ability to accumulate both total biomass (fresh and dry) per unit of stem height and foliar biomass (fresh and dry) per unit of leaf area (Figure 6C-F), while lines 1-3 had a 22 and 19% reduced ability to accumulate tot dry biomass per unit of stem height and dry foliar biomass per unit of leaf area, respectively (Figure 6D and F). Line 1-3 showed also an 8% reduction in chlorophyll content, whereas line 4-6 had the same chlorophyll content compared to wild-type plants (Figure 6G). Although a certain level of phenotypic variation was observed in transplastomic plants, these results demonstrate that the heterologous production of these diterpenes is not associated with any aboveground phenotypic penalties in selected lines 4-6. Despite this phenotypic variation, all PdTPS-CbGGPPS T/L lines (1-6) were not statistically different for their ability to produce tubers (Figure 6H and I) compared to wild-type control plants. Specifically, both the total fresh weight of tubers (Figure 6H), and the total number of tubers (Figure 6I) per plant were not statistically different compared to controls, demonstrating intact total yield per plant and tuber size in transplastomic plants producing diterpenes.

**Figure 6:**
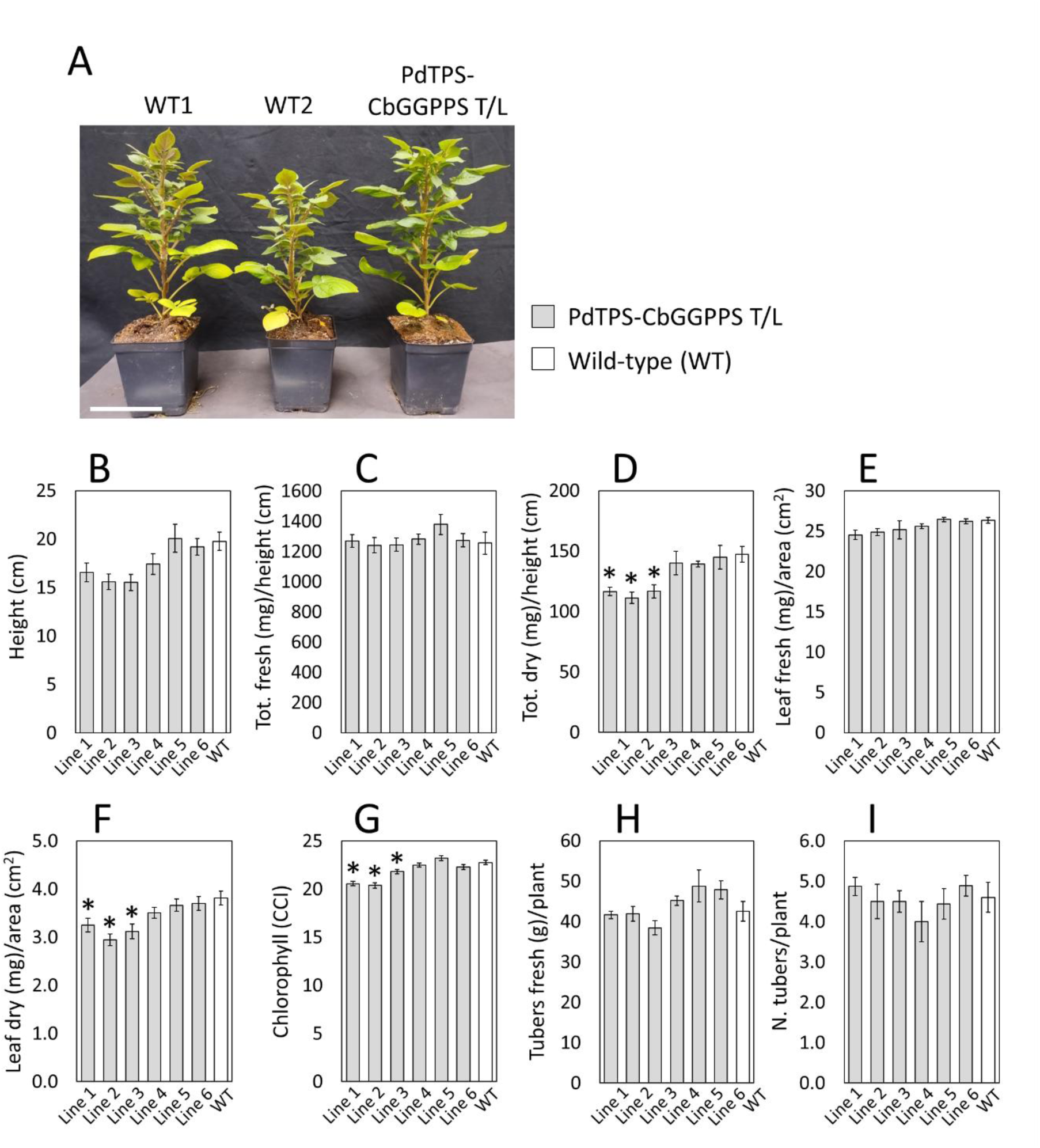
Growth characteristics of PdTPS-CbGGPPS T/L transplastomic lines. **(A)** Images showing PdTPS-CbGGPPS T/L lines (1-6) at anthesis (∼7-8-week-old) along with wild-type (WT1 and 2) control plants grown on pot in a growth chamber. Scale bar: 10 cm. Various phenotypic plant characteristics are shown: **(B)** Plant height (cm); **(C)** ratio of total fresh weight (mg) to plant height (cm); **(D)** ratio of total dry weight (mg) to plant height (cm); **(E)** ratio of leaf fresh weight (mg) to foliar area (cm^2^); **(F)** ratio of leaf dry weight (mg) to foliar area (cm^2^); **(G)** chlorophyll content index (CCI); **(H)** total fresh weight of tubers (g) per plant; **(I)** total number of tubers per plant. Data are expressed as mean ± standard error (SE) of 8 plants per each transplastomic line and wild-type plants. Means were compared using ANOVA Post-Hoc Dunnett’s t test (p<0.05, C, E-G, and I) or Post-Hoc Dunnett’s T3 (p<0.05, B, D, and H) and statistically significant lower means compared to the wild-type reference group are indicated (*).

## DISCUSSION

This study demonstrates that the chloroplast genome can serve as an effective platform for stable, high-value diterpene biosynthesis in an agronomically relevant crop, establishing potato as a tractable system for plastome-based engineering of the terpene pathway beyond the *Nicotiana* model. The identification of the *trnT/trnL* locus as an optimal integration site, combined with the strategic co-expression of a geranylgeranyl diphosphate synthase (CbGGPPS) to balance precursor supply, resolved the growth penalties initially associated with transplastomic lines and unlocked substantially elevated diterpene accumulation (Figure 5 and 6).

The success of this work is grounded in a deep understanding of the evolutionary history of terpene pathways across the plant kingdom. Of the two parallel terpene biosynthetic pathways in the plant cell, the MVA pathway was inherited from the eukaryotic host ancestor and the MEP pathway from the endosymbiotic cyanobacterium, following primary endosymbiosis ^60–62^. Although the genes encoding the MEP pathway were entirely transferred from the cyanobacterial genome to the nuclear genome over evolutionary time, all MEP pathway enzymes are re-imported into the chloroplast, enabling the pathway continues to operate within this organelle ^63^. This evolutionary history directly points to the chloroplast as a logical and tractable compartment for engineering terpene pathways. Through the MEP pathway, one key short-chain isoprenyl diphosphate precursor is produced: GGPP. GGPP synthesis, catalyzed by GGPP synthase (GGPPS), a member of the IDS family, is broadly conserved across the plant kingdom ^53^. Notably, a recent study on the evolution of IDS family revealed that the GGPPS gene originated in cyanobacteria was retained in the chloroplast genome in glaucophytes, the earliest-diverging lineage of the plant kingdom, whereas in all other plant lineages, including red algae, green algae, and land plants, GGPPS has been relocated to the nuclear genome ^38^. Reintroducing the GGPPS gene into the chloroplast genome therefore represents a restoration of the ancestral evolutionary design.

Our design of terpene pathway engineering was further inspired by current understanding of the evolutionary history of TPS genes. TPS genes are specific to land plants within the plant kingdom, and the ancestral TPS has been hypothesized to be a tridomain (α, β, and γ domains) bifunctional diterpene synthase capable of catalyzing sequential class II and class I reactions ^18,39,42,43^. This hypothesis was significantly strengthened by the discovery of structurally analogous tridomain bifunctional TPS genes in bacteria, supporting the idea that plant TPSs were originally acquired from a bacterial ancestor via horizontal gene transfer ^64,65^. Strikingly, tridomain bifunctional TPS enzymes are conserved across all lineages of non-flowering land plants, including lycophytes, ferns, and gymnosperms, but are entirely absent in flowering plants ^43^. Over the course of plant evolution, one ancestral bifunctional enzyme appears to have been split into two distinct monofunctional diterpene synthases, one retaining class I activity and the other class II activity, which now operate in tandem ^66^. This evolutionary transition toward monofunctionality in flowering plants conferred a significant advantage: the decoupling of the two catalytic activities created new opportunities for combinatorial pairing of class I and class II enzymes, substantially amplifying the chemical diversity of the diterpene repertoire. However, this functional separation comes at a cost when metabolic specificity and pathway efficiency are the primary objectives, as the requirement for coordinated expression of two separate enzymes introduces additional complexity and potential bottlenecks. Using a bifunctional diterpene synthase from a lycophyte (PdTPS) therefore represents a deliberate return to ancestral enzyme architecture, streamlining biosynthesis into a single catalytic step while maintaining product specificity.

A critical consideration in engineering diterpene biosynthesis in the chloroplast is the central role of GGPP in primary metabolism. Beyond serving as the substrate for diterpene secondary metabolites, GGPP is an essential branching point for multiple core physiological processes, supplying precursors for gibberellins, chlorophylls, tocopherols, plastoquinone, and carotenoids ^55–57^. Given this broad metabolic centrality, it is unsurprising that redirecting a significant portion of the chloroplastic GGPP pool toward the production of secondary diterpenes imposes measurable costs on plant physiology. In our transplastomic lines expressing the bifunctional diterpene synthase alone, the most conspicuous phenotypic consequence was a bushy, dwarfed growth habit (Figure 3 and Figure S4), a phenotype consistent with gibberellin deficiency resulting from competition for the shared GGPP substrate ^67–69^. The co-expression of a GGPP synthase within the chloroplast genome effectively resolved this metabolic imbalance by augmenting the local supply of GGPP (Figure 5 and 6), thereby alleviating competition between the endogenous primary metabolic demands ^55–57^ and the introduced diterpene biosynthetic pathway. This result underscores the importance of precursor engineering as an integral component of any metabolic engineering strategy targeting GGPP-derived products and demonstrates that pathway flux and plant fitness need not be mutually exclusive objectives when the supply-demand balance is appropriately managed.

Preserving normal plant growth is essential for the potential use of this genetic engineering platform for terpene engineering in crops for improving important agricultural traits. For this purpose, we considered both key genetic engineering factors to enable efficient metabolic engineering and genetic tools to minimize impacts on plant phenotype due to transgene integration and overexpression. A key aspect of this study was selecting an optimal integration site into a ∼0.7 kb intergenic *trnT*/*trnL* region of plastome that minimizes negative effects on endogenous proximal genes of this integration site (Figure 1). Notably, in PdTPS-CbGGPPS T/L plants, vector integration and overexpression of three transgenes (*aadA*, *PdTPS* and *CbGGPPS*) exciding by several folds the level of the endogenous *rbcL* gene (Figure 5) did not alter the expression levels of *trnT*/*trnL* flanking genes (Figure S5), supporting optimal vector design and location in the plastome. In addition to being suitable for transgene overexpression using the regulatory elements (promoter and UTRs) indicated in Figure 5, this *trnT*/*trnL* integration site is located in the large single copy (LSC) region of the plastome, which allows full homoplasmy to be achieved within one vegetative generation in tissue culture. It should be noted that integration sites located in the inverted repeat (IRA and IRB) regions of potato plastome typically require two vegetative generations of embryogenic *calli* induction ^36^, increasing the time required to produce transgenic plants. Furthermore, the utilization of integration sites that allow full homoplasmy in a single vegetative generation will also be essential for redesigning this *trnT*/*trnL* vector into a marker-free mFree vector ^49^, enabling the production of homoplasmic marker-free lines with complete removal of the selectable marker gene and further accelerating the path to commercialization of novel transplastomic potato varieties in agriculture.

This study demonstrates for the first time that the plastome of a major crop plant can be successfully harnessed for high-value diterpene production. Beyond the specific case of potato, the principles established here, including rational design of optimal transformation vectors, precursor supply optimization, and evolutionarily informed enzyme architecture, provide a generalizable framework extensible to other crops and terpene classes. The chloroplast offers compelling but largely untapped advantages for crop biotechnology, including high-level transgene expression, maternal inheritance minimizing gene flow risk, and a stromal environment naturally rich in MEP pathway intermediates. Looking forward, stacking additional pathway genes within the plastome could enable complete biosynthetic routes to terpenes of biological and industrial relevance. Combined with computational pathway modeling and directed enzyme evolution, this work repositions chloroplast genome engineering not as a niche tool confined to model systems, but as a practical strategy for the sustainable, contained, and scalable production of plant natural products in agriculture.

## MATERIALS AND METHODS

### Plant genotype and growth experiments

In this study, *Solanum tuberosum* L. var. Desirée (potato) was used. Magenta GA7 vessels (W × L × H = 77 mm × 77 mm × 97 mm) containing MS Reg medium (4.33 g/L Murashige and Skoog (MS) basal salt mixture; 25 g/L sucrose; 100 mg/L myo-inositol; 170 mg/L sodium phosphate monobasic monohydrate; 440 mg/L calcium chloride dihydrate; 0.9 mg/L thiamine-HCl; 2 mg/L glycine; 0.5 mg/L nicotinic acid; 0.5 mg/L pyridoxine-HCl; 1X MS vitamins; 3 g/L phytagel; pH 5.8) were used for routine *S. tuberosum* propagation in tissue culture, while transplastomic plants were propagated in the same vessels containing MS Reg medium supplemented with 200 mg/L of spectinomycin.

For phenotypic studies, transplastomic and wild-type plants were grown in ∼1.3 L pots containing Pro-Mix BK25 (Griffin Greenhouse Supplies). Plants were grown until flowering (anthesis) in controlled environmental facilities (growth chamber or greenhouse) using a light/dark cycle of 16/8 hours, a light intensity of ∼350-400 μE/m^2^/s, and a temperature of 22-24°C. At the end of the growth experiment, several phenotypic parameters were collected, including the plant height, total plant and leaf fresh and dry biomass, leaf surface area, and leaf chlorophyll content (chlorophyll content index, CCI). For dry biomass determination, plant tissue was kept for ∼1 week at 60°C until completely dried. The leaf surface was collected using a scanner, and the area was calculated using the ImageJ software (National Institutes of Health, NIH). The leaf chlorophyll content was obtained using a portable CCM-200 plus chlorophyll meter (OPTI-SCIENCES). Phenotypic data were processed using Microsoft Excel software (Microsoft) and mean separation was evaluated using ANOVA Dunnett’s t or T3 (p<0.05) using the IBM SPSS software (IBM).

### Construction of chloroplast transformation vectors

For construction of the chloroplast T/L transformation vector backbone (pUC T/L), two 1.5 kb-long *trnT* (from nt 46617 to nt 48116) and *trnL* (from nt 48117 to nt 49616) homologous arms from *S. tuberosum* plastome (GenBank: DQ386163.2) including *Bpi*I restriction sites (TGCC/GGGA) for Golden-Gate cloning, were inserted into an in-house produced pUC plasmid ^48,49^. For the construction of GFP T/L vector, the *mEmerald* (*Prrn-T7G10-mEmerald-psbA-3’UTR*) and *aadA* (*PpsbA-5’UTR-aadA-rbcL-3’UTR*) expression cassettes were assembled using modular cloning kits as previously described ^70,71^ and cloned into the *Bpi*I site of the pUC T/L plasmid backbone. The two genes of the terpene pathway, *PdTPS* (GenBank: OL989442.1) ^18^ and *CbGGPPS* (NCBI Ref: YP_009504633.1) ^38^ used to design the PdTPS T/L and PdTPS-CbGGPPS T/L vectors, were gene synthetized (Integrated DNA Technologies, IDT) as lvl0 modules for Golden Gate cloning ^71^. For construction of PdTPS T/L and PdTPS-CbGGPPS vectors, the *aadA* (*Prrn-RBS-aadA-psaC-3’UTR*), *PdTPS* (*PpsbA-5’UTR-PdTPS-petD-3’UTR*), and *CbGGPPS* (*PrbcL-RBS-CbGGPPS-psbA-3’UTR*) expression cassettes were assembled using modular cloning kits ^70,71^ and cloned into *Bpi*I sites of the pUC T/L plasmid backbone. The *aadA* and *PdTPS* expression cassettes were used to design the PdTPS T/L vector, while the *aadA*, *PdTPS*, and *CbGGPPS* expression cassettes were used for PdTPS-CbGGPPS T/L vector design. The MoClo Toolkit used in the procedures of cloning was a gift from Sylvestre Marillonnet (Addgene kit # 1000000044) ^71,72^, while the MoChlo (Modular Cloning Chloroplast Toolbox) was a gift from Scott Lenaghan (Addgene kit #1000000156) ^70^.

### Production of potato transplastomic plants in tissue culture

For production of transplastomic plants, leaf tissue from the top of the canopy of ∼4-week-old potato plants grown in tissue culture was used. For each transformation event using the gene-gun PDS-1000/He delivery system (Bio-Rad), a total of ∼6 cm^2^ of leaf tissue placed on the surface of M6M medium (4.33 g/L MS basal salt mixture; 1x Gamborg B5 vitamins; 30 g/L sucrose; 18.2 g/L mannitol; 18.2 g/L sorbitol; 0.8 mg/L zeatin riboside (ZR); 2 mg/L 2,4-dichlorophenoxyacetic acid (2,4-D); 3 g/L phytagel; pH 5.8) was used. For each transformation, 0.3 mg of 0.6 µm gold particles (Bio-Rad) coated with 1 µg of purified plasmid DNA were bombarded into leaf tissue positioned at 6 cm-distance from the gun, using 1,100 psi rupture disks (Bio-Rad), as previously described ^36^. Embryogenic green calli were induced from transformed leaf tissue after ∼8-12 weeks in tissue culture, incubating tissue in M6 medium (4.33 g/L MS basal salt mixture; 1x Gamborg B5 vitamins; 30 g/L sucrose; 0.8 mg/L zeatin riboside (ZR); 2 mg/L 2,4-dichlorophenoxyacetic acid (2,4-D); 400 mg/L spectinomycin; 3 g/L phytagel; pH 5.8) for the first ∼4 weeks and then Ti medium (4.33 g/L MS basal salt mixture; 1x Gamborg B5 vitamins; 16 g/L glucose; 3 mg/L zeatin riboside (ZR); 2 mg/L indole acetic acid (IAA); 1 mg/L gibberellic acid (GA_3_); 400 mg/L spectinomycin; 3 g/L phytagel; pH 5.8) for the following ∼4-8 weeks, as described before ^36^. Embryogenic green calli were kept proliferating for ∼4 weeks in DH medium (2.16 g/L MS basal salt mixture NH_4_NO_3_^-^ free; 268 mg/L NH_4_Cl; 1x Nitsch vitamin mixture; 2.5 g/L sucrose; 36.4 g/L mannitol; 100 mg/L casein hydrolysate; 80 mg adenine hemisulfate; 2.5 mg/L zeatin riboside (ZR); 0.1 mg/L indole acetic acid (IAA); 400 mg/L spectinomycin; 3 g/L phytagel; pH 5.8) and thereafter, the production of transplastomic plants was induced from calli incubated in MON medium (4.33 g/L MS basal salt mixture; 1x Gamborg B5 vitamins; 30 g/L sucrose; 0.1 mg/L naphthaleneacetic acid (NAA); 5 mg/L zeatin riboside (ZR); 400 mg/L spectinomycin; 3 g/L phytagel; pH 5.8) after ∼4 weeks, as described previously ^36^. Transplastomic plants were propagated in tissue culture in Magenta GA7 vessels containing MS Reg medium supplemented with 200 mg/L spectinomycin.

### Total DNA extraction and Southern blot analysis

Total genomic DNA was extracted from potato leaf tissue using the CTAB method as described previously ^36^. For this purpose, ∼50 mg of leaf tissue was pulverized in liquid nitrogen and resuspended into 500 μL of CTAB buffer (2% hexadecyltrimethyl ammonium bromide; 1% (w/v) polyvinyl pyrrolidone; 100 mM Tris-HCl; 1.4 M NaCl; 20 mM EDTA; 0.1 mg/ml RNaseA). Homogenized tissue was incubated for 30 minutes at 60°C, thereafter, genomic DNA was extracted in chloroform/isoamyl alcohol (24:1), precipitated, washed, and resuspended in ∼50 µL sterile milli-Q water as previously described ^36^.

For Southern blot analysis, a ∼0.5 kb digoxigenin-labelled probe against the *trnT*/*trnL* integration site (*rps4* gene) was synthetized using the PCR DIG Probe Synthesis Kit (Roche) and the pair of primers 1Fw/1Rv following the manufacturer’s instructions. For each sample, a total amount of 1 µg of genomic DNA was subjected to restriction digestion using the indicated restriction enzymes in the corresponding result sections and separated by electrophoresis on a 0.9% agarose gel. After separation, the gel was depurinated, denatured and transferred to Hybond-N+ nylon membrane (GE Healthcare) as previously described ^36^. The DIG-labelled probe was detected using the anti-digoxigenin-AP Fab fragments detection kit (Roche) following the manufacturer’s instructions.

### Total RNA extraction, cDNA synthesis and qRT-PCR

The CTAB method was also used for preparation of high-quality total RNA from leaf tissue as described previously ^73^. For each extraction, ∼50-100 mg of leaf tissue was ground in liquid nitrogen and resuspended into 500 μL of RNase-free CTAB buffer supplemented with 286 mM β-mercaptoethanol. Samples were incubated for 30 minutes at 60°C, and then, total RNA was extracted in chloroform/isoamyl alcohol (24:1), precipitated, washed, and resuspended in ∼50 µL sterile RNase-free ultrapure water (Zymo Research). Total RNA preparations were subjected to DNase treatment (DNase I-XT, New England Biolabs) to remove any DNA contamination and then purified on column using the Monarch® Spin RNA Cleanup Kit (New England Biolabs) following the manufacturer’s instructions. The cDNA synthesis was performed using the ZymoScript RT PreMix Kit (Zymo Research) following the manufacturer’s instructions. Reactions of qRT-PCR were prepared in 15 µL volume using 1X PowerUp™ SYBR™ Green Master Mix (Thermo Fisher Scientific), using cDNA samples dilute 1:3 and 0.5 µM of each primer. The *aadA* (GenBank: ARK38551.1), *PdTPS* (GenBank: OL989442.1) and *CbGGPPS* (NCBI Ref: YP_009504633.1) genes along with the endogenous reference gene *rbcL* (Gene ID: 4099985) were amplified using primer pairs 2Fw/2Rv, 3Fw/3Rv, 4Fw/4Rv, and 5Fw/5Rv, respectively (Table S1). PCR amplification data were collected using a QuantStudio 6 Flex Real-Time PCR System (Thermo Fisher Scientific) following the manufacturer’s instructions. The relative transgene expression data versus the *rbcL* reference gene of the plastome were expressed using the equation 2^-ΔCt^. Mean separation was evaluated using ANOVA Dunnett’s T3 (p < 0.05) using the IBM SPSS software for statistical analysis (IBM).

### Terpene synthase enzyme assays

The protein expression construct based on the pEXP5-CT/TOPO vector harboring *PdTPS* gene has been previously described ^18^. The construct was transformed into *E. coli* BL21 CodonPlus cells (Invitrogen) for protein expression. *E. coli* cells harboring the expression construct were grown at 37°C until reaching an OD_600_ of 0.6. Gene expression was induced by adding isopropyl β-D-1-thiogalactopyranoside (IPTG) to a final concentration of 1 mM. Protein isolation was performed following the method described by Zhuang et al ^74^. The recombinant enzyme assay was performed in a 2 mL GC vial containing 50 μL of bacterial extract, 50 μL of 10 mM MgCl_2_, and 10 μM GGPP. A solid-phase microextraction (SPME) fiber coated with 100 μm polydimethylsiloxane was inserted into the headspace of the vial. After 30 min incubation at 25°C, the SPME fiber was directly injected into the injection port of a Shimadzu GC-2010 plus coupled to a Shimadzu QP2010 SE quadrupole mass selective detector with splitless mode for product identification. Mass spectra were acquired using electron ionization (EI) at 70 eV with the ion source maintained at 200°C. Helium served as the carrier gas at a constant flow rate of 1 mL min^-1^. Compound separation was achieved using a low-polarity Shimadzu SH-I-5MS capillary column (30 m × 0.25 mm i.d., 0.25 μm film thickness). The oven temperature program started at 60°C with a 6 min hold, followed by an increase to 300°C at a rate of 5°C min^−1^, and a final hold at 300°C for 5 min.

### Diterpene identification using GC-MS

Potato leaves were ground into a fine powder with liquid nitrogen, and 0.05 g of ground tissue was transferred into a 2 mL GC vial. 200 μL of 20% sodium chloride solution was added and mixed thoroughly. A SPME fiber was inserted into the headspace of the vial for volatile collection for 30 min at 25°C. The SPME fiber was subsequently injected into the GC–MS system for analysis. The GC–MS conditions were the same as those described for terpene synthase enzyme assays.

## Supporting information

This file includes all supplementary Figures and Tables of this manuscript

## Acknowledgements

This research was funded by the University of Tennessee, startup funds and seed grants allocated to A.O., and an AgResearch SPRINT award from the University of Tennessee, Institute of Agriculture allocated to F.C. and A.O. This work was also supported by USDA Hatch Grants to A.O. and F.C.

## Author contributions

A.O. and F.C. conceived, designed and supervised this study; A.O., S.A.M. and G.K. produced transplastomic plants; S.A.M., M.M., I.A.F.Q. and A.O. performed molecular and phenotypic characterization of transplastomic plants; X.C. performed TP enzymatic assays and GC-MS experiments and processed data. A.O., F.C. and X.C. interpreted data; A.O., F.C. and X.C. wrote the first draft of the manuscript; everyone edited and approved the final version of the manuscript.

## Competing interests

The authors declare no competing interests.

## Supplementary information

This file includes all supplementary Figures and Tables listed below:

Figure S1. TPS phylogenetic tree.

Figure S2. Mass spectrums of terpenes.

#Figure S3. Expression of genes of the *trnT*/*trnL* integration site in PdTPS plants.

Figure S4. Growth characteristics of PdTPS T/L lines *in vitro*.

Figure S5. Expression of genes of the *trnT*/*trnL* integration site in PdTPS-CbGGPPS plants.

Table S1. Primers used in this study.

## Notes

### Competing Interest Statement

The authors have declared no competing interest.

