## Supplementary material for "Chloroplast genome engineering of potato enables diterpene production without agronomic penalty": This file includes all supplementary Figures and Tables of this manuscript

This file includes all supplementary Figures and Tables listed below:

Figure S1. TPS phylogenetic tree.
Figure S2. Mass spectrums of terpenes.
Figure S3. Expression of genes of the *trnT*/*trnL* integration site in PdTPS plants.
Figure S4. Growth characteristics of PdTPS T/L lines *in vitro*.
Figure S5. Expression of genes of the *trnT*/*trnL* integration site in PdTPS-CbGGPPS plants.
Table S1. Primers used in this study.

**Figure S1**

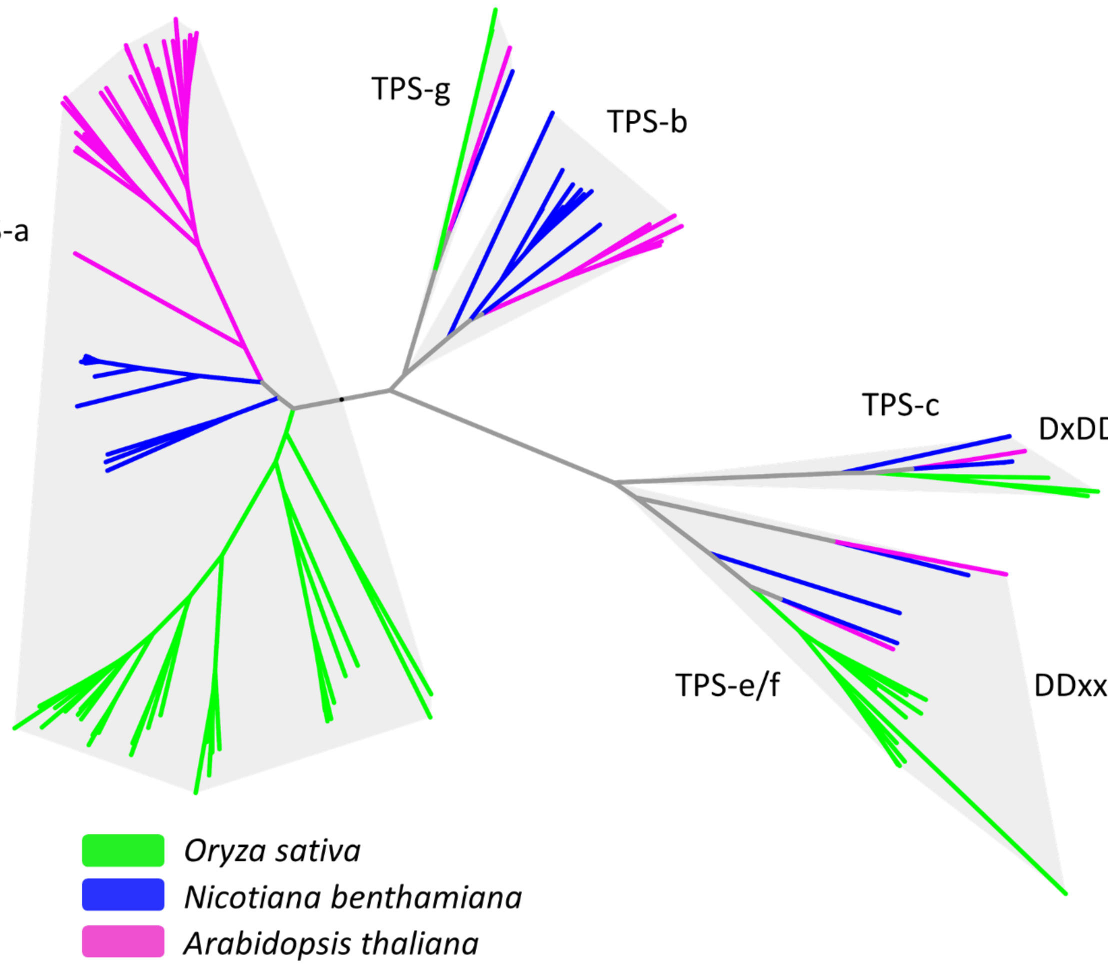


**Figure S1. TPS phylogenetic tree.** Phylogenetic tree of TPSs from *Nicotiana benthamiana* together with TPSs from *Arabidopsis thaliana*, and *Oryza sativa*. Moel predicted by Prottest is JTT+G+F. Members of the TPS-c subfamily contains only the 'DxDD' motif while members of the TPS-e/f subfamily contain only the 'DDxxD' motif.

**Figure S2**

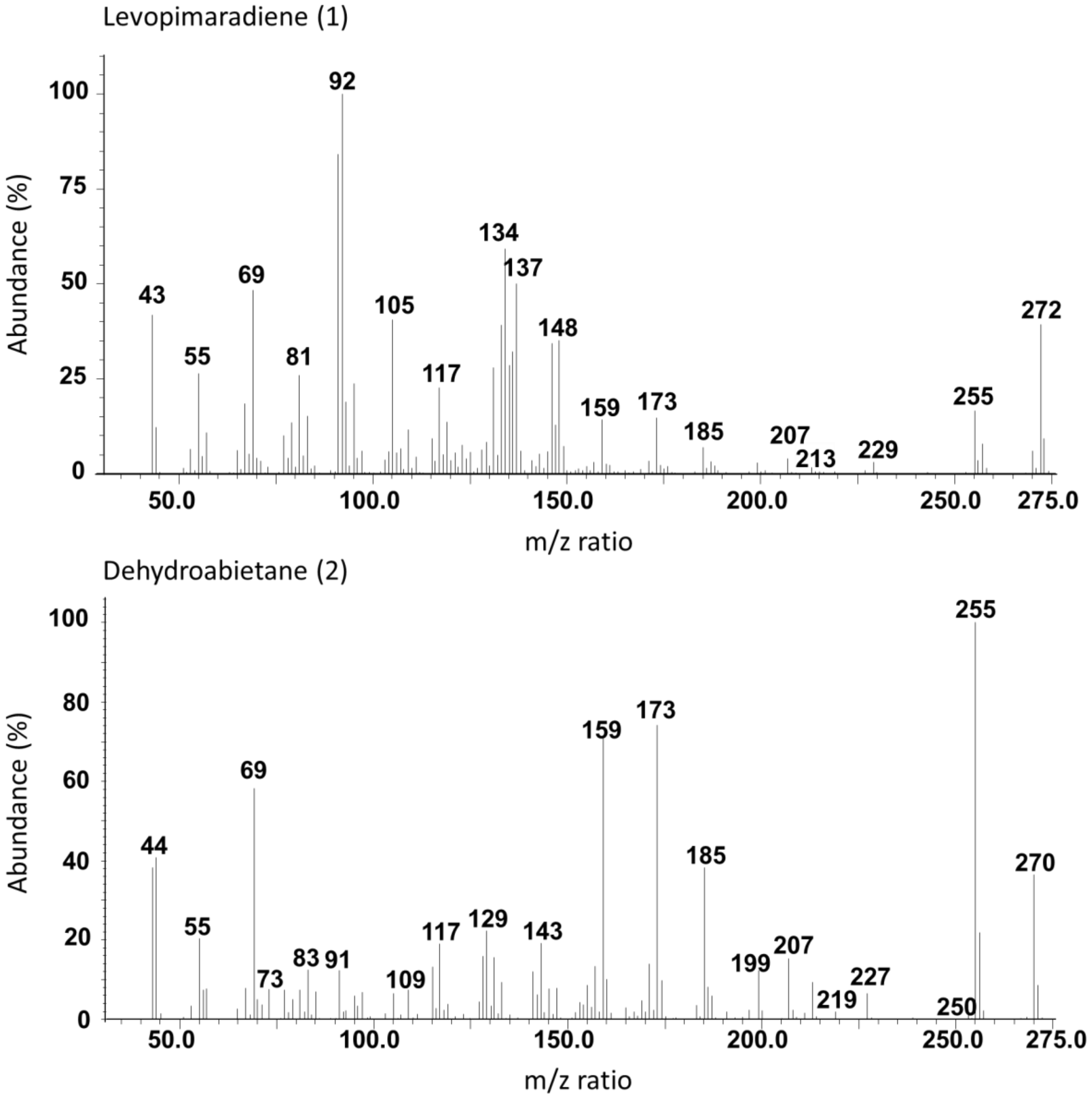


**Figure S2. Mass spectrums of terpenes.** Mass spectrums of levopimaradiene (1) and dehydroabietane (2) from PdTPS enzymatic assay are indicated. The relative abundance (%) and the m/z ratio are indicated in the y and x axis, respectively.

**Figure S3**

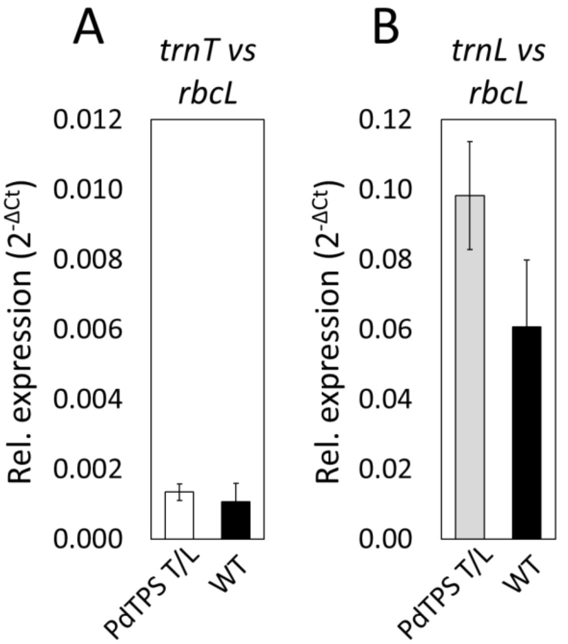


**Figure S3. Expression of genes of the *trnT/trnL* integration site in PdTPS plants. (A-B)** qRT-PCRs were performed using cDNA samples from PdTPS T/L and wild-type plants, and primers for amplification of *trnT* (A) and *trnL* (B) located in proximity of the integration site. Graphs represent the relative expression ($2^{-\Delta Ct}$) of either *trnT* or *trnL* versus the *rbcL* gene indicated as the average of all PdTPS T/L lines (1-3). Data represent mean ± SE (standard error) of 3 biological replicates per each line and 3 technical replicates per each biological. Mean separation was evaluated using t test ($p<0.05$).

**Figure S4**

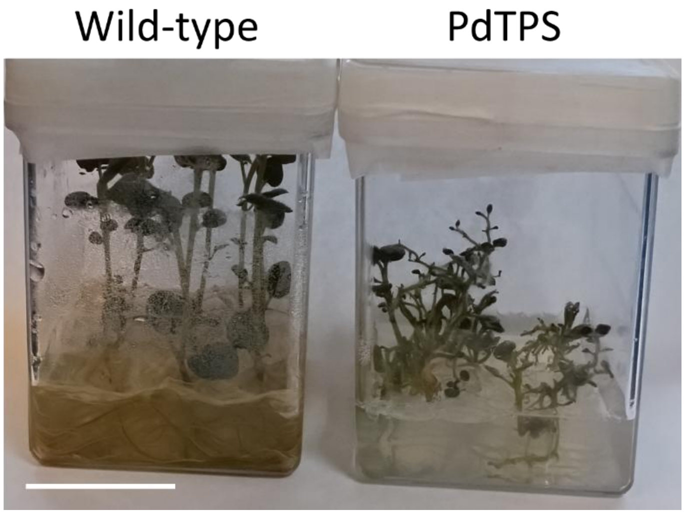


**Figure S4. Growth characteristics of PdTPS T/L lines *in vitro*.** Four-week-old wild-type potato along with PdTPS T/L line 1 grown in tissue culture are shown. Scale bars: 5 cm.

**Figure S5**

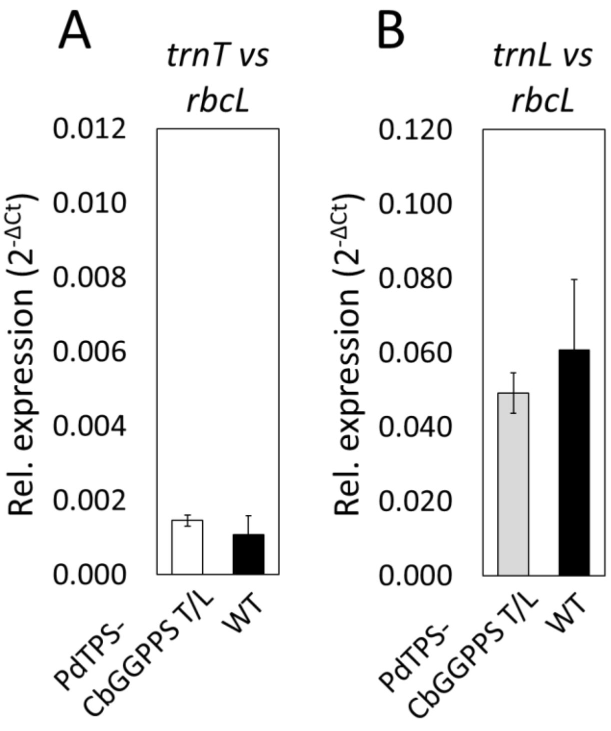


**Figure S5. Expression of genes of the *trnT*/*trnL* integration site in PdTPS-CbGGPPS plants. (A-B)** qRT-PCRs were performed using cDNA samples from PdTPS-CbGGPPS T/L and wild-type plants, and primers for amplification of *trnT* (A) and *trnL* (B) located in proximity of the integration site. Graphs represent the relative expression ($2^{-\Delta Ct}$) of either *trnT* or *trnL* versus the *rbcL* gene indicated as the average of all PdTPS-CbGGPPS T/L lines (1-6). Data represent mean ± SE (standard error) of 3 biological replicates per each line and 3 technical replicates per each biological. Mean separation was evaluated using t test ($p<0.05$).

**Table S1. Primers used in this study.** The id, the name and the 5'-3' nucleotide sequence are indicated for each primer. Primers are subdivided into forward (Fw) and reverse (Rv).

**Forward primers**

| **id** | **name** | **sequence (5'-3')** |
|---|---|---|
| 1 Fw | rps4-probe Fw | GGGGTTTGCAGCGATAACTCGGTAT |
| 2 Fw | SmR-q Fw | TGAGGCGCTAAATGAAACCT |
| 3 Fw | PdTPS-q Fw | GCAGACATGGAGTGACCGAA |
| 4 Fw | CbGGPPS-q Fw | TGGAAATGGCTTTACCAACAGC |
| 5 Fw | rbcL-q Fw | AGATCTGCGAATCCCTGTTG |

**Reverse primers**

| **id** | **name** | **sequence (5'-3')** |
|---|---|---|
| 1 Rv | rps4-probe Rv | CAGTAAACTTCGCTTCATTTAGTTCAG |
| 2 Rv | SmR-q Rv | TACTGCGCTGTACCAAATGC |
| 3 Rv | PdTPS-q Rv | TGCGAGGTAAGCTTTGGAGT |
| 4 Rv | CbGGPPS-q Rv | GTTTTCCGCGTCGATAAGAATC |
| 5 Rv | rbcL-q Rv | CAGGGGACGACCATACTTGT |
